# Off-target integron activity leads to rapid plasmid compensatory evolution in response to antibiotic selection pressure

**DOI:** 10.1101/2022.04.29.490043

**Authors:** Célia Souque, José A. Escudero, R.Craig MacLean

## Abstract

Integrons are mobile genetic elements that have played an important role in the dissemination of antibiotic resistance. Under stress, the integron can generate combinatorial variation in resistance cassette expression by cassette re-shuffling, accelerating the evolution of resistance. However, the flexibility of the integron integrase site recognition motif hints at potential off-target effects of the integrase on the rest of the genome that may have important evolutionary consequences. Here we test this hypothesis by selecting for increased piperacillin resistance populations of *P*.*aeruginosa* with a mobile integron containing a hard-to-mobilise beta-lactamase cassette to minimize the potential for adaptive cassette re-shuffling. We found that integron activity can both decrease overall survival rate but also improve the fitness of the surviving populations. Off-target inversions mediated by the integron accelerated plasmid adaptation by disrupting costly conjugative genes otherwise mutated in control populations lacking a functional integrase. Plasmids containing integron-mediated inversions were associated with lower plasmid costs and higher stability than plasmids carrying mutations, albeit at a cost of reduced conjugative ability. These findings highlight the potential for integrons to create structural variation that can drive bacterial evolution, and they provide an interesting example showing how antibiotic pressure can drive the loss of conjugative genes.

## Introduction

Mobile integrons are genetic shuffling devices heavily involved in the spread of antibiotic resistance (Escudero et al., 2015). They consist of an integrase gene followed by an array of promoterless genes cassettes (Stokes and Hall, 1989), predominantly encoding for resistances genes (Partridge et al., 2009). Cassettes are expressed from a promoter located at the end of the integrase gene, such that their expression level is dependent on their position within the array (Collis and Hall, 1995; Souque et al., 2021). When bacteria are exposed to antibiotics, the generation of DNA damage activates the SOS response and the expression of the integron integrase (Guerin et al., 2009). The integrase enzyme then leads to cassette re-shuffling, duplication and deletion, generating combinatorial variation in both cassette presence/absence and expression levels, modulating bacteria antibiotic resistance levels (Barraud and Ploy, 2015; Souque et al., 2021).

This cassette shuffling is possible through the recognition by the integrase of two motifs found within the integron: the double-stranded *attI* site (Hansson et al., 1997), located at the start of the array, and the single stranded *attC* sites (Francia et al., 1999), located at the end of the cassettes. Recombination between two *attC* sites lead to the excision of the intervening cassette (Collis and Hall, 1992), while the *attC x attI* reaction promotes integration of the cassette at the start of the array (Collis et al., 1993). Integrase recognition of these binding sites requires little sequence homology, relying instead on short degenerated core sequences (for the *attI* sites) (Hall et al., 2000) or the recognition of two to three extrahelical bases acting as structural landmarks (*attC* sites) (Bouvier et al., 2009). This flexibility creates the potential for the integron to generate ‘off-target’ recombination with sites outside of the integron, such as the insertion of integron cassette in other parts of the genome (Recchia and Hall, 1995; Segal and Elisha, 2006). These off-targets effects of the integrase are likely to be deleterious: they may lead to the formation of chromosomal dimers that have to be resolved before cell division, or potentially create genomic instability through the deletion of entire genomic regions (Harms et al., 2013). These reactions may therefore compromise the evolutionary benefits associated with integron cassette re-arrangements. Moreover, these shuffling evolutionary benefits are likely to be dependent on the presence of highly mobile cassettes with strong positional effect on expression levels. As integron cassette mobility is highly variable, with cassette recombination frequency spanning several order order of magnitudes depending on their *attC* site and their position within the array (Aubert et al., 2012; Loot et al., 2010; Nivina et al., 2016), the nature of the integron cassette cargo is likely to play a key role in shaping the relative importance of on-target and off-target effects of the integron.

Our previous work investigating the evolutionary benefits of integrase activity used a model system involving a highly mobile *aadB* cassette that that was under strong selection for an optimal position (Souque et al., 2021). Here we test the evolutionary benefits of integron activity using *blaVEB-*1, a β-lactam resistance cassette that has low mobility and weak positional effects on antibiotic resistance. Under these conditions, the adaptive value of the cassette re-shuffling is likely to be weak, and side-effects of integrase activity are expected to play a greater role in determining the evolutionary impact of integron activity. In this scenario we observed integrase activity decreased overall population survival but had a beneficial impact on the fitness of the surviving populations through the generation of extensive plasmid backbone rearrangements, creating novel evolutionary pathways.

## Results

### The impact of cassette position varies between cassettes

We used a combinatorial three-cassette class 1 integron system consisting of all six possible configurations of three resistance cassettes, *dfrA5, aadB*, and *blaVEB-1*, each encoding resistance to a different antibiotic family (trimethoprim, aminoglycoside and beta-lactams) (Souque et al., 2021) and measured the impact of cassette position on resistance levels. While previous results showed a strong impact with the *aadB* cassette of cassette position resistance on gentamicin (Souque et al., 2021), the impact of position was much more subtle (Figure 1a) or inexistent (MIC equal or above 1500mg/L for all array) when looking at piperacillin and trimethoprim resistance and the *blaVEB-1* and *dfrA5* cassette respectively. At the transcription level, as expected, cassette position had an overall clear impact on transcript levels. However, and perhaps surprisingly, the impact of cassette position on transcription was variable depending on the cassettes and was weaker for *dfrA5* and *blaVEB-1* when compared to the *aadB* cassette(Supplementary Figure S1). Adding to the previous observations that the gentamicin resistance gradient provided by the *aadB* cassette was complemented by a decrease in translation, these results show mobile integron cassettes expression and the resistance levels they provide are the product of interactions between transcription, translation, and the nature of the resistance mechanism itself.

**Figure 1.**
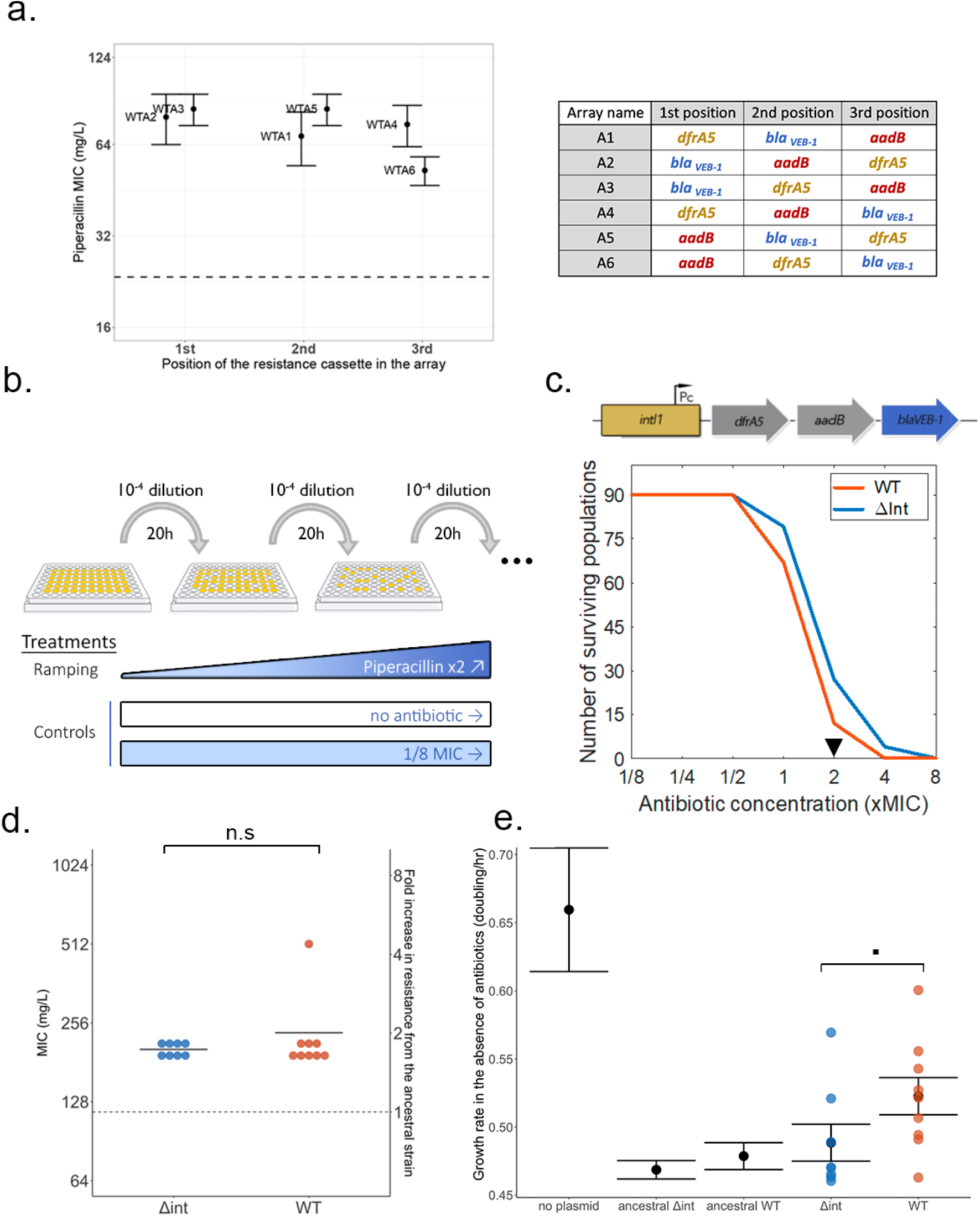
Integrase activity impacts bacterial evolution under antibiotic selection without altering resistance phenotypes. A. impact of *blaVEB-1* cassette position on resistance levels (piperacillin minimum inhibitory concentration) using a combinatorial integron array system (right). Error bars represent the standard error of three biological replicates. The dotted line represents *P. aeruginosa* resistance level in the absence of the plasmid. B. ‘evolutionary ramp’ experimental design. C. Top: representation of the WTA4 array, with the beta-lactamase cassette *blaVEB-1* highlighted in blue. Bottom: Survival rates of the WT and *Δint* populations over time. The black triangle represents the time-point selected for further phenotypic and genomic analysis. D. Final piperacillin resistance level of evolved populations sampled from the x2 MIC time point. Each dot represents the average of three independent biological replicates. The dotted line corresponds to the resistance level of the ancestral strains. Difference in resistance between the two genotypes was compared using a Fisher t-test (t = -0.91247, df = 8.214, p-value = 0.3875) E. Final growth rate in the absence of antibiotic of the evolved populations. Each dot represents the average of three independent biological replicates, with the dotted line corresponding to the growth rate of the ancestral strains. Error bars represents standard error. Difference in growth rate in the evolved populations between the two genotypes was compared using a Wilcoxon rank sum test (W = 16, p-value = 0.059).

The ‘adaptation on demand’ model of the integron as an evolutionary catalyst assumes that antibiotic pressures impose strong selection for resistance cassettes to be positioned in first position, as we have reported for *aadB*. However, these results show that cassette position can have weak effects on resistance levels provided by cassettes such as *blaVEB-1* and *dfrA5*.

### Integrase activity impact bacterial evolution under antibiotic selection without altering resistance phenotypes

Given the varying impact of cassette position on resistance levels, we aimed to test the impact of the integron on resistance evolution when selection for cassette re-arrangement is weak. Starting from the array WTA4, which contains the *blaVEB-1* cassette in last position, we built a mutant with a dysfunctional integrase *Δint*A4 but providing similar initial levels of resistance. Using an ‘evolutionary ramp’ design (Bell and MacLean, 2018), we compared the ability of both strains to evolve resistance to increasing concentrations of the antibiotic piperacillin (Figure 1B). We passaged 90 independent populations of each strain for 7 days in doubling concentrations of antibiotics, starting from 1/8 MIC and ending at 8 times the original MIC. As control, 15 populations of each strain were passaged with either low but constant selection (constant piperacillin concentration at 1/8 MIC) or in the absence of selection (no antibiotic).

We followed survival rates through OD_600_ readings and observed resistance evolution (survival at x1 MIC and above) in most but not all populations, with extinction of all populations at eight times the original MIC (Figure 2C). Surprisingly, we observed a better survival rate of the *Δint* populations, with some populations surviving up to x2 MIC. While minor, the difference in survival rates was statistically significant (log-rank test: Chisq= 12.4 on 1 degree of freedom, p= 4e-04).

**Figure 2:**
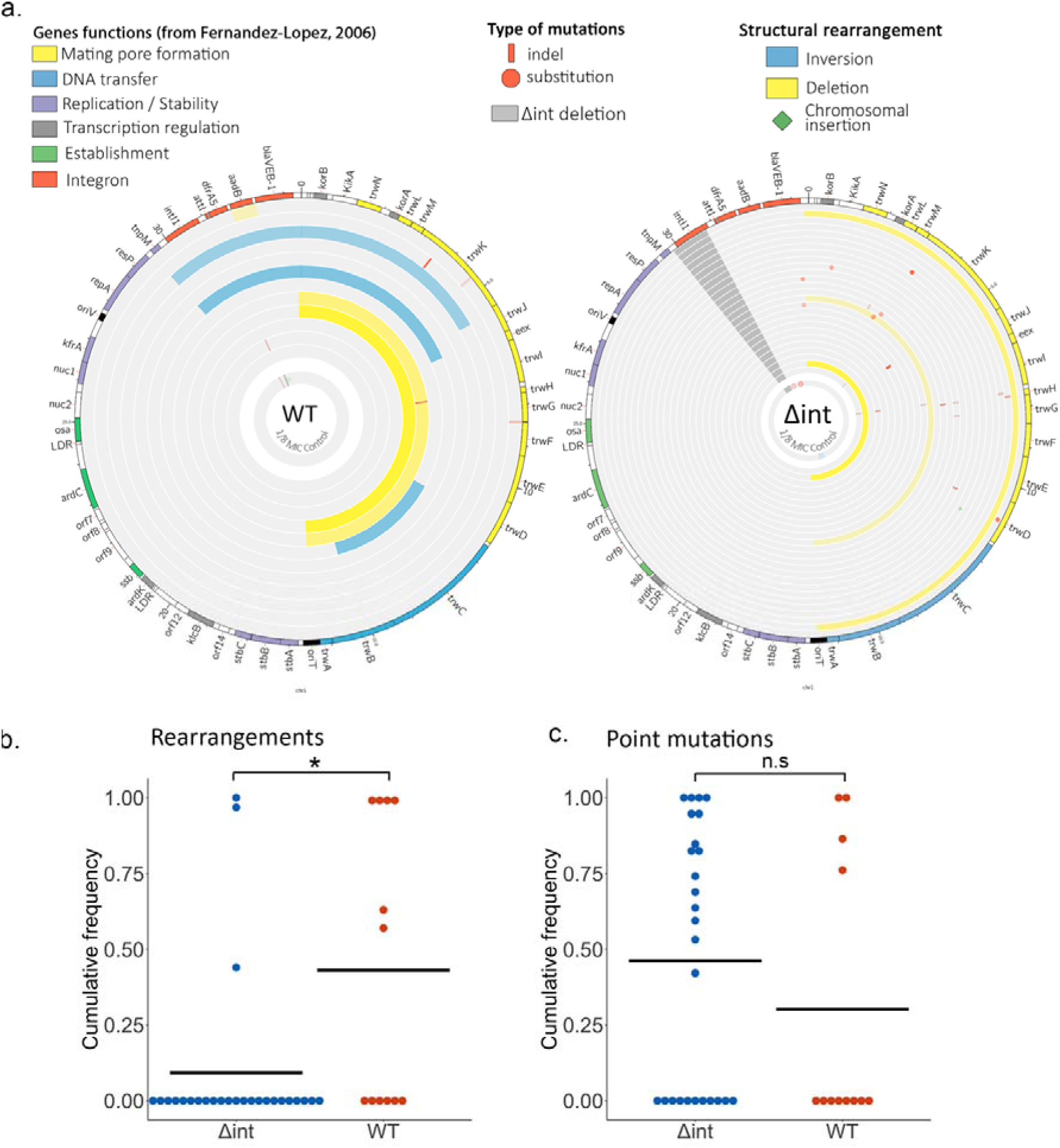
Integron activity shapes plasmid evolution through extensive structural rearrangements. A. Representation of the plasmid mutations and rearrangements in the surviving WT (left – 12 populations) and *Δint* (right – 26 populations) populations at x2 MIC, mapped to the R388 reference sequence. Each circle represents a separate population, with the inner circle representing the variants present in an equimolar pool of six 1/8 MIC control populations. Large scale inversions and deletions are represented in blue and yellow respectively, while indels and single nucleotide substitutions are represented in red. The color intensity represents the frequency of the corresponding mutation. The dark grey area in the *Δint* populations represents the location of the *IntI1* deletion. As DNA was extracted from entire populations, symbols may overlap when both rearrangements and mutations are identified in separate sub-populations. The function of each R388 gene as described in (Fernández-López et al., 2006) is indicated by a specific color in the outer circle. B Cumulative frequency per population and genotype of large scale rearrangements (inversion, deletion and cassette rearrangement) – Frequencies compared using the Wilcoxon rank sum test (W = 94.5, p-value = 0.010) C. Cumulative frequency per population and genotype of point mutations (SNP and short indels) - Frequencies compared using the Wilcoxon rank sum test Wilcoxon rank sum test (W = 183, p-value = 0.37)

To test the impact on integrase activity on the evolution of resistance, we measured both piperacillin resistance and fitness in the absence of antibiotic for a subset of WT and *Δint* evolved populations that evolved to 2x MIC (Figure 2D and 2E). We observed no difference in final piperacillin resistance between the two genotypes, suggesting that the integrase had no effect on the evolution of resistance *per se*. However, we were able to identify a small increase in final fitness in populations with a functional integrase, suggesting that the integrase facilitates the evolution of low-cost resistance and/or compensatory evolution (Wilcoxon rank sum test, W = 16, p-value = 0.059).

### Integron activity shapes plasmid evolution through extensive structural rearrangements

We then investigated the genotype of the evolved ramping populations at x2 MIC using whole genome sequencing. Focusing first on the integron-bearing plasmid, we observed striking mutations and structural rearrangements targeting the conjugation machinery of the R388 plasmid backbone, with 58% (7 out of 12) of the WT and 61% (16 out of 26) of the *Δ*int populations presenting mutations and/or rearrangements in the *trw* operon (Figure 2a). Alongside the 16.6kb long deletion of the DNA transfer replication and mating pore formation modules of the conjugation machinery, we also identified three large scale inversions in this region of the plasmid: two inversions from *resP* to *trwJ* and one from *trwD* to *trwB (*respectively 9.5kb, 10.1kb and 4.3kb in size). Mutations were found in the *trwK, trwH, trwD, trwG* and *trwM* genes, with 12 out of 17 mutations being either non-sense or frame-shifting mutations. Finally, we identified the insertion within the *trwD* gene of one *Δ*int population of a 1.2kbp IS element initially present on the chromosome.

By comparison, the integron array and its beta-lactam cassette were much more conserved. We observed one sole instance of cassette arrangement, the excision of the *aadB* cassette, in one WT population. Mutations in the integron array were also sparse, as we observed mutations and/or indels in only two *Δint* populations (a non-synonymous SNP within *blaVEB-1*), and in one WT population (a 20bp deletion between the *attI* site and the start of the *dfrA5* cassette).

While the *Δint* and WT populations evolved through modifications of similar targets, we observed a striking difference in the way these alterations were achieved: the conjugation machinery was disabled in *Δ*int populations mainly through mutations, while WT populations were enriched in structural rearrangements, with half of the WT populations presenting some type of rearrangement (Fig 2B). While the extensive 16kb deletions were found in both WT and *Δint* populations, the large-scale inversions were also specific to the populations with a functional integrase.

Surprisingly, we observed a different pattern, both in mechanism and in target, in the control populations evolved at constant concentration of piperacillin. The pooled controls contained predominantly mutations targeting the integron array and not the conjugation machinery, both in the WT and *Δint* populations. We identified four nonsense and frameshifting mutations targeting the *dfrA5* cassette, two mutations in the intergenic region between the *aadB* and *blaVEB-1* cassettes, one insertion within *aadB* of an *ISPa11* (Partridge and Hall, 2003) initially located on the chromosome, and a 131bp deletion within the integrase of the WT control. The only rearrangement observed was a short 773bp inversion within *trwC* in the *Δint* control. The comparison between variants observed in the control and piperacillin selected populations suggests that antibiotic pressure generated selection for the loss of conjugation machinery, whereas selection drove the loss of costly resistance genes in the absence of increasing antibiotic pressure.

Finally, the plasmid was completely lost at this time point in the pooled controls passaged without antibiotic while plasmid copy number remained similar between the ancestral and evolved populations of both genotype in the ramping and constant antibiotic treatment (Figure S2).

While we observed an impact of integrase activity on plasmid evolution, no differences were identifiable on the chromosome (Supplementary table S1 & Supplementary Figure S3). WT and *Δint* populations had an average cumulative mutation frequency on the chromosome of 0.78 and 0.66 respectively, and no extensive deletion or inversion could be identified. Parallel evolution was observed in the *PA1766-PA1768* operon, whose genes are predicted to be involved post translation modification of cell wall proteins (Gutu et al., 2013) and a *PA1767* mutant was shown to provide a moderate increase in piperacillin resistance (Dötsch et al., 2009). Other mutated genes were shown to be linked with antibiotic resistance, such as *galU* (Dean and Goldberg, 2002; Sanz-García et al., 2018), *PA1195* (Dötsch et al., 2009), *clpA* (Jorth et al., 2017; Sanz-García et al., 2018), and *infB* (Sanz-García et al., 2018), but no link could be found for the remaining 16 out of 26 targeted genes. Excluding one mutation, all these mutations were either non-synonymous or intergenic and therefore still show sign of selection, and we speculate that some of these mutations may be compensatory mutations to offset the cost of plasmid carriage. For example, we identified mutations in *hslU* and *clpA*, two ATP-binding protease, and *rne*, a ribonuclease, which have been shown to interact with plasmids replication proteins (Kubik et al., 2012; Söderbom et al., 1997; Wickner et al., 1994), a driver of plasmid cost in *P. aeruginosa* (San Millan et al., 2015).

### Plasmids inversion junction sites correspond to integrase secondary sites

As plasmid backbone rearrangements, and especially inversions, were found more frequently in *WT* than in *Δint* populations, we looked at the junctions sequence for signs of integrase activity (Figure 3). The recognition characteristics of *intI1* require little sequence identity, with only a three base long conserved motif, 5’-GNT-3’, which can be extended to form 7bp GTTRRRY motif (Hall et al., 1991), and where cleavage usually happens between the base A and C of the complementary strand (Frumerie et al., 2010).

**Figure 3:**
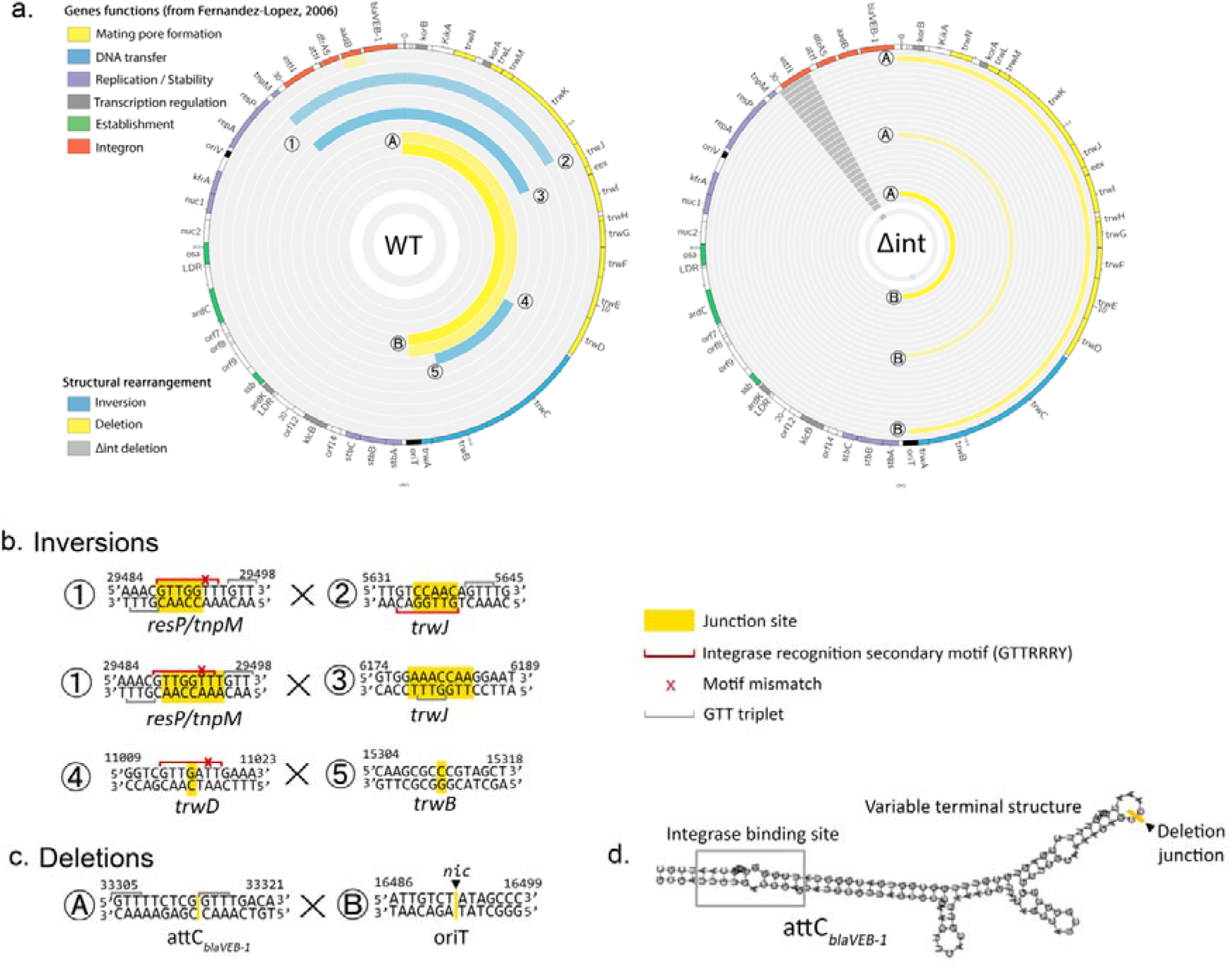
Plasmids inversion junction sites correspond to integrase secondary sites. a) Plasmids rearrangements junctions labelled 1 to 4 (inversions) and A to B (deletions). b & c) Junction site of each inversion and deletion, based on the reference sequence. The crossover junction site is indicated in yellow. When the crossover point is unclear due to sequence homology between each junction the entire homology is highlighted. Motifs close to the GTTRRRY integron secondary sequence are indicated by a red line, with red crosses indicating the potential mismatch. Simple 5’-GTT-3’ triplets are indicated by a grey line. The *nic* site of the origin of transfer *oriT* is indicated by a black triangle. d.) folding of *attC*_*blaVEB-1*_ calculated using ViennaRNA (Lorenz et al., 2011), with the deletion junction in the variable terminal structure represented in yellow.

The inversions between *resP/tnpM* and *trwJ* have a strong signal of integrase activity: while the precise cross-over point cannot be determined due to sequence homology a 5’-GTT-3’ triplet, often extendable in a motif close to GTTRRRY, can be identified in both sides of the inversion. Even more strikingly, the motifs on each side of the inversion are in inverted orientation from each other, with one located on the top strand and another on the bottom strand. It has been shown that the integrase can create sequence inversions through intramolecular *attI×attI* reactions when the sites are in inverted orientation (Escudero et al., 2016) In this case the Holliday junction formed during the first step of the recombination process is resolved through second strand exchange instead of replication, as is the case with the canonical *attI×attC* or *attC×attC* reactions.

The involvement of the integrase in the *trwD×trwB* inversion is less straightforward. We identified a motif close to GTTRRRY at the *trwD* end, but with a cleavage point after the 5’-GTT-3’ triplet. It has been shown that the constraints on the second nucleotide of the triplet are fairly lax (Frumerie et al., 2010; Hansson et al., 1997), and that cleaving can happen between the G and the A of a 5’-GAT-3’, as observed here, especially in the case of reaction with secondary sites (Francia et al., 1993). However, we were not able to identify any 5’-GTT-3’ triplet in the vicinity of the *trwB j*unction. As a control, we also investigated the small inversion located within the 1/8 MIC *Δint* pooled sample: we identified one 5’-GTT-3’ triplet on one side, but not on the other, and no motif close to GTTRRRY.

In contrast, extended deletions found in both WT and *Δint* populations share the same boundary site which does not show integrase binding motifs. One boundary is found in *attC*_*blaVEB-1*_ and its variable terminal structure, a part of the *attC* site usually not involved in integrase recognition. The other end of deletion is located precisely at the *nic* site of the R388 *oriT*, where the *trwC* relaxase binds during conjugation and initiates the nicking of the DNA to create single stranded DNA, which is then exported to the recipient cell and recircularized. It has been shown that *trwC* can mediate site-specific recombination between two *oriT* sites even in the absence of conjugation (Llosa et al., 1994) which leads to loss of the intervening DNA (Draper et al., 2005). It is unclear if the *trwC* enzyme can bind to the sequence at the *attC*_*blaVEB-1*_ junction site: recognition of the *nic* site is supposed to be highly specific (Lucas et al., 2010) and no sequence homology could be seen between *oriT* and *attC*_*blaVEB-1*_. As the limiting step for *oriT* recombination is the generation of a single strand at *oriT* to allow *trwC* nicking (César et al., 2006), recombination between the two sites may be facilitated by the *attC* site unique secondary structure which make the junction site available as single stranded DNA.

### Integron activity alleviates plasmid fitness cost and improves plasmid persistence through loss of the conjugation machinery

A key insight from sequencing the evolved populations is that integrase containing populations tend to evolve by structural re-arrangements of the R388 conjugative apparatus, whereas *Δint* populations evolve via points mutations in conjugative genes. We then investigated if the higher proportion of structural rearrangements in populations with a functional integrase could explain the higher final fitness observed previously. To test this idea, we extracted the plasmids out of subset of WT and *Δint* evolved populations and transformed them into the ancestral chromosomal background. We found that the ancestral R388 plasmid imposes a large fitness cost and this cost was ameliorated by compensatory evolution during the piperacillin selection experiment (Figure 4a). Crucially, plasmids with extensive rearrangements of the conjugation machinery imposed less of a fitness cost than the unevolved plasmids or plasmids with point mutations. In contrast, both mutations and rearrangements provided similar level of resistance to piperacillin (Figure 4b).

**Figure 4:**
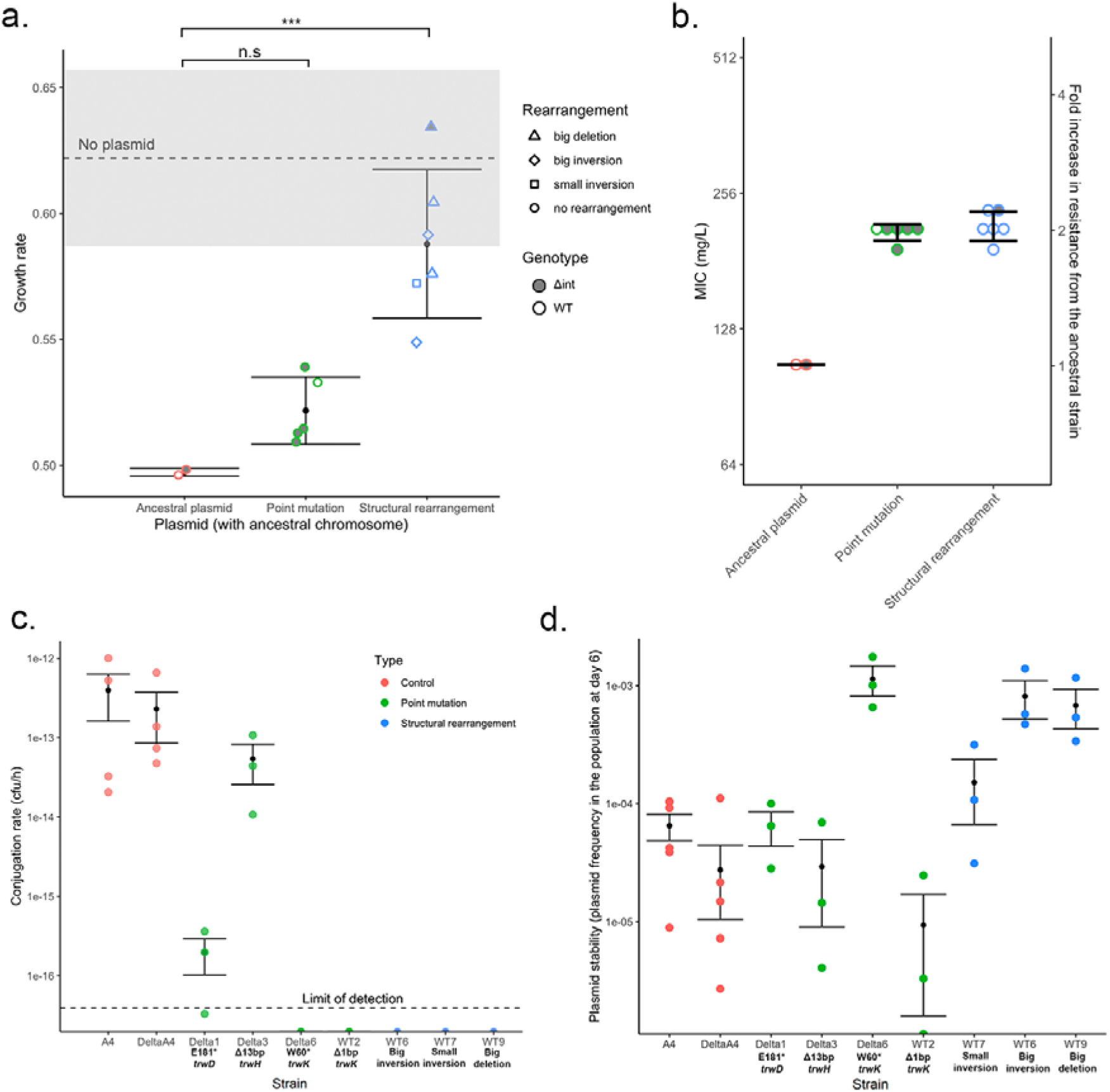
Loss of the conjugation machinery increases piperacillin resistance and improves plasmid persistence. a&b. Growth rate in the absence of antibiotic (a) and Piperacillin MIC (b) of the evolved plasmids when transferred back in the ancestral chromosome. Each dot represents the average of three independent biological replicates. The error bars correspond to the standard deviation with each category. The greyed area represents the standard error of the growth rate in the absence of plasmid. (c). Conjugation rate estimation for a selection of mutations and rearrangements after filter mating. The dotted line corresponds to the average sensitivity limit of the assay. (d) Plasmid stability estimated by the final plasmid frequency obtained after transferring 3 to 5 independent populations of each strain in the absence of antibiotic for 6 days. Error bar represents standard error.

The strong compensatory effects associated with structural re-arrangements suggests that these variants increased fitness by eliminating the conjugative ability of the R388 plasmid. To test this hypothesis, we measured the conjugation rate of evolved plasmids. Both inversions and deletions were enough to stop the plasmid from conjugating, while mutations often provided an intermediate phenotype (Figure 4c).

Both selection and horizontal transfer impact the ability of plasmids to persist in bacterial populations. The observed structural re-arrangements decrease the fitness costs of plasmids carriage, which should improve the ability of the plasmid to persist through vertical transmission. However, these re-arrangements also reduce conjugative ability, hindering the ability of R388 to persist via horizontal transfer. To understand the net effect of plasmid evolution on stability, we transferred independent populations of each plasmids in the absence of antibiotic during six days, and measured the ratio of cells containing the plasmid at the final time point. Rearranged plasmids were present at a ten-fold higher frequency than ancestral or mutated plasmids, showing these rearrangements improved plasmid persistence (Figure 4d).

## Discussion

The ‘adaptation on demand’ model shows the clear benefits of the integron when cassettes have high mobility and cassette position is under strong selection. However, it is important to emphasize that integron cassettes display a diversity of fitness costs, recombination rates and positional effects, all of which may impact the benefits of integrase activity. Using the beta-lactamase encoding cassette *blaVEB-1*, which has been shown to have reduced mobility and a weak cassette expression gradient, we investigated the evolutionary impact of integrase activity in an experiment where selection for cassette re-arrangement is weak. Under these conditions integrase expression decreased the survival rate of the populations under antibiotic treatment, revealing a cost to integrase activity (Fig 1c). However, the integrase had a positive impact on the fitness of the surviving populations by generating inversions in conjugative genes that effectively compensated for the high costs associated with the R388 plasmid (Fig 2a). Integrase-mediated compensatory adaptation increased the stability of the R388 plasmid in the absence of antibiotic pressure, albeit at a cost of decreased conjugative ability (Fig 4d). These results highlight the limitation of the adaptation on demand model, and they emphasize the potential importance of off-target effects to moderate the fitness costs and benefits of the integron.

Our study showed the potential benefits of integrase activity are not limited to the canonical *attI x attC* and *attC x attI* reactions, but can also lead to adaptive inversions between secondary sites through *attI x attI*-like reactions. The mechanism behind the reactions between *attI* sites had been studied in detail (Escudero et al., 2016), but its potential adaptive benefits have remained elusive. *AttI-*like sites can be found frequently across genomes (Harms et al., 2013) and these adaptive rearrangements between secondary sites may be common. Integrons consisting only of an integrase and without cassettes arrays (so called *In0*) have been found several time across bacteria genomes (Cury et al., 2016; Néron et al., 2022), and adaptive benefits of off target integrase activity may help to maintain these cassette-less integrons.

While the inversions we observed had clear signs of integrase activity, the mechanism behind the extensive *attC x oriT* deletions we observed is more elusive. The relaxase *trwC* has been shown to mediate site-specific excision between two *oriT* sites and is triggered by the generation of single stranded DNA at one of the *oriT* site (César et al., 2006). The unique single stranded folding of *attC* sites may make them unexpected hotspots for recombination through their variable terminal structure (VTS). The ubiquity of integron cassettes on plasmids suggests that integron cassette and their *attC* sites may play a role in plasmid remodelling even in bacteria containing only dysfunctional integron integrase pseudogenes (Nemergut et al., 2008).

When comparing these results to our previous study, it is noteworthy that with the *aadB* and *blaVEB-1* cassettes, the intensity of the cassette expression gradient depending on position in the array was positively correlated with mobility. Depending on the resistance mechanism they encode, not all cassettes may be able to generate a significant resistance gradient in the typically 1 to 6 cassettes length of a mobile integron (see *dfrA5*). Similarly higher expression levels which could be provided by cassette duplication may be deleterious, as is the case for some beta-lactamase cassettes (Rajer and Sandegren, 2022). In that case, and especially if the cassette has a high fitness cost, a highly mobile *attC* site would have little benefits to the cassette but would be linked to a higher rate of cassette excision, leading to the selection of *attC* sites with lower excision rate. Integron cassettes as mobile genetic elements may be subject to their own level of selection, leading to traits which may not be constantly beneficial for their bacterial host or the integrase itself.

Finally, this study showed a striking parallel evolution of conjugation deficient plasmids in response to antibiotic selection pressure. R388 was found to be a very costly plasmid in *P. aeruginosa*, but its fitness cost could be greatly improved by disabling the conjugation machinery. It is not surprising that conjugative genes carry a fitness cost as they are often associated with an increased cellular burden (Porse et al., 2016; San Millan and MacLean, 2017). In *P. aeruginosa*, it has been shown that the conjugative pilus causes a membrane perturbation that triggers the ‘tit-for-tat’ response and activates the T6SS (Ho et al., 2013) while conjugative genes can also be targeted by CRISPR-Cas systems (Wheatley and MacLean, 2021). Not all conjugative plasmids are costly *in P. aeruginosa*, with the plasmid paKD1 containing tightly repressed conjugation genes actually increasing PA01 fitness (San Millan et al., 2018) but a badly regulated conjugation machinery could generate a cascade of costly reactions. Surprisingly, we found that disrupting the conjugative machinery led to an unexpected increase in piperacillin resistance, revealing a novel cost of conjugation independently on its impact on *P. aeruginosa* fitness. We can speculate that the presence of conjugative pili on the bacteria membrane may make them more susceptible to beta-lactams antibiotics. This effect is relatively subtle (2x change in MIC), but it clearly led to the accelerate loss of conjugative genes in the presence of antibiotics. The generality of this mechanism is unclear, but our results raise the tantalizing possibility that antibiotic conjugation-induced susceptibility may be an unexplored selection pressure driving the acquisition and maintenance of beta-lactam resistance genes on conjugative plasmids.

## Material and methods

### Strains and growth conditions

The strains and plasmids used in this paper are listed in Supplementary table S2. Unless stated differently, bacteria were grown in LB Miller broth at 37*C with shaking (225 rpm) and plasmid maintenance was guaranteed through the addition of 100 mg/L of ceftazidime.

### Minimum inhibitory concentration determination

Minimum inhibitory concentrations (MIC) for each antibiotic were determined following the broth microdilution method from the Clinical and Laboratory Standards Institute (CLSI) guidelines. 5×10^5^ c.f.u bacteria inocula were prepared in cation-adjusted Mueller-Hinton Broth 2 (MH2) using individual colonies grown on selective agar and incubated in doubling concentrations of antibiotics for 20h in three technical replicates. Cultures optical density (OD_600_) was then read using a Biotek Synergy plate reader and wells were considered empty when the overall OD600 was under 0.1. The MIC for each assay was defined as the minimal concentration in which growth was inhibited in all three technical replicates. The final MICs values are the average of two to three replicate assays (from separately prepared inocula, on different days).

### Growth rate determination

Growth rates in the absence of antibiotics were determined using a Biotek Synergy plate reader. Populations were inoculated in similar conditions to the MIC assay (5×10^5^ c.f.u bacteria inocula prepared in cation-adjusted (MH2) from individual colonies) and followed for 24h at 37*C with periodic shaking, with OD_600_ measurements taken every 15 minutes. Intrinstic growth rate were determined using the Growthcurver package (Sprouffske and Wagner, 2016). Each measurement was done in three technical replicates (separate wells from the same inocula) and replicated three times over different days (biological replicates).

### Experimental evolution

#### Minimum inhibitory concentration in experimental conditions

Minimum inhibitory concentrations for piperacillin were determined for all strains in conditions matching the experimental evolution set-up: overnight cultures were diluted 1/10 000 and supplemented with doubling concentrations of piperacillin. MICs were determined after 20h of incubation. This process was repeated twice. In these conditions the MIC were 64 mg/L of piperacillin for both the PA01:WTA4 and PA01:*Δint*A4 strains.

#### Evolutionary ramp experiment

90 populations of each strain (PA01:WTA4 and PA01:*Δint*A4) were initially inoculated from individual colonies grown on selective agar in MH2 at a piperacillin concentration of 1/8 MIC. The outer wells of every plate were kept as negative controls and strain distribution among the plates was kept balanced to limit plate effects. Plates were kept at 37°C with shaking at 225 rpm. Every day the populations were diluted 1/10 000 in a doubling concentration of MIC until reaching x8 MIC. As controls, 15 populations of each strain were transferred either without antibiotic or at a constant piperacillin concentration of 1/8 MIC. The OD of each population was measured each day and a population was considered extinct when its OD fell below 0.1 after 20h of incubation. All populations were frozen in 15% glycerol every two days.

### Genome analysis

#### DNA extraction

PA01:WTA4 and PA01:Δ*int*A4 populations from the x2 MIC time-point were re-grown from frozen stock in LB Miller media supplemented with a piperacillin concentration of x1 MIC (ramping populations), x1/8 MIC (1/8 MIC controls), no antibiotic (no antibiotic controls) or 100 mg/L of ceftazidime (ancestral PA01:WTA4 and PA01:Δ*int*A4 populations). DNA extraction was performed using the DNeasy Blood & Tissue Kit (Qiagen) through the QiaCube extraction platform. Samples were treated with RNAse before extraction and DNA concentration was measured using the Quantifluor dsDNA system (Promega). For each control treatment (1/8 MIC and no antibiotic, for each genotype) the DNA of six populations was pooled in equimolar ratio and sequenced as one sample.

#### Next Generation Sequencing

Library preparation and sequencing was performed by the Oxford Genomics Centre at the Wellcome Centre for Human Genetics. Sequencing was performed in two batches. Twelve samples of each genotype as well as the ancestral populations (PA01, PA01:WTA4 and PA01:Δ*int*A4) and the pooled controls were first sequenced on the NextSeq 500 sequencing system (Illumina). Five PA01:WTA4 populations as well as the ancestral strains were contaminated by phages during the DNA extraction process and were re-extracted and re-sequenced alongside the remaining PA01:Δ*int*A4 populations (14 populations) on the HiSeq4000 platform.

#### Bioinformatic pipeline

PCR duplicates and optical artifacts were removed using MarkDuplicates (Picard toolkit) while low quality bases and adaptors were trimmed using Trimmomatic v.039 (Bolger et al., 2014) and overall quality control was performed using FastQC (Andrews *et al*., 2012) and multiQC (Ewels et al., 2016). Phage contamination in the first batch of samples was determined at this stage.

Variant calling and rearrangement identifications were performed using the *breseq* pipeline in polymorphism mode (Barrick et al., 2014). Variants present in the un-evolved ancestor populations were filtered out irrespective of frequency. Due to the lower quality of the samples processed by NextSeq 500, which led to an over-identification of low frequency variants, a filtering process harsher our previous analysis (Souque et al., 2021) was carried out: any variant present in the controls evolved without antibiotic or at a frequency of less than 30% was removed from the ramping populations dataset. The evidence for any variant above this threshold and present in several NextSeq 500 populations was manually examined and the variant removed if present in a region of low read quality. For the pooled 1/8 MIC populations a lower threshold of 5% (which corresponds to a variant present in 30% of one out of 6 populations) was applied and manual examination was carried out to exclude low quality variants. Additional evidence for plasmid rearrangements identified by *breseq* (large scale deletions, inversions, chromosomal insertion) was obtained using plasmidSPAdes (Antipov et al., 2016).

#### PCR analysis

Cassette rearrangement and structural variants were also confirmed by PCR. Surviving populations at the x2MIC time point were plated using a pin replicator on LB agar supplemented with piperacillin at a concentration corresponding to x1 MIC and incubated for 48h at 37°C. 12 populations of each genotype were picked randomly, boiled for 10 minutes at 95°C in distilled water and screened for cassette rearrangement. The cassette rearrangement panel consisted of 4 reactions: amplification of the integrase size (as control for cross-contamination), localisation of the *blaVEB-1* cassette relative to the start of the integrase, localisation of the *blaVEB-1* cassette relative to the end of the *dfrA5* cassette and identification of potential *blaVEB-1* cassettes duplications (primer sequences are indicated in Table S3). These PCRs were performed using the GoTaq G2 DNA mastermix (Promega) for 30 cycles with 30s at 95°C denaturation, 30s at 55°C annealing and 3 minutes at 72°C elongation.

The inversions and deletions identified through whole genome sequencing (inversions between *resP* x *trwJ* and *trwD × trwB*, deletions between *blaVEB-1* and *stbA)* were confirmed through a second PCR panel. Inversions were detected using a pair of primers specific to the integrase and *trwG* gene, or the t*rwD* and *trwB* genes, and binding on the same strand, therefore only producing an amplicon if the intervening sequence is reversed. Deletions were targeted using primers specific to *blaVEB-1* and *stbA*: given *blaVEB-1* and *stbA* are separated by 17 kbp on the ancestral plasmid, amplification is only possible in the presence of deletion bringing the primers closer together. The following PCR protocol was used: 30s at 95°C denaturation, 30s at 55°C annealing and 1 minute at 72°C elongation.

### Evolved plasmids analysis

#### Transformation of the evolved plasmids into the ancestral chromosomal background

Extraction of the plasmids from the evolved populations was performed using the QIAprep Miniprep (Qiagen) on the QiaCube extraction platform from liquid culture from the frozen populations grown in LB Miller supplemented with 32mg/L of piperacillin. Extracted plasmids were then electroporated back into PA01 (Choi and Schweizer, 2006). Presence of the plasmid was confirmed by PCR. Colony PCR and, when required, Sanger sequencing of the amplicons was performed to identify single colonies containing either the rearrangements or the selected mutations.

#### Conjugation rate determination

Determination of the conjugation rates of the ancestral and evolved plasmids was performed using filter mating with a rifampicin resistant PA01 acceptor (D512G *rpoB* mutant from (Qi et al., 2014)). Populations of donor and recipients were grown from overnight for 3h30 with (donors) and without (recipient) antibiotics, washed and resuspended in NaCl (0.9%). Donor and recipient populations were mixed at a 1:1 ratio, deposited on filter on agar plate without antibiotic alongside pure donor/recipients control and incubated for 1h. Populations were then resuspended in NaCl 0.9%. Concentrations of donors, recipients and transconjugants were determined at both T0 and Tf using appropriate antibiotics (ceftazidime 100mg/L, rifampicin 64mg/L and both combined). Conjugation rate was calculated using the Simonsen method (Simonsen et al., 1990).

#### Plasmid maintenance over time

Populations with ancestral and evolved plasmids were inoculated from single colonies in the presence of antibiotic and passaged for 6 days in the absence of antibiotic with a similar dilution (1:10 000) as the experimental evolution experiment. After 6 days the ratio of plasmid containing cells was determined by serial dilution plating with (for the cfu of the plasmid containing cells) and without ceftazidime (for the cfu of the entire population). The presence of horizontal gene transfer in these conditions was confirmed by mixing a subset of the populations with rifampicin resistant bacteria after day 1 and plating them after 20h on rifampicin + ceftazidime plates.

#### Statistical analysis

Statistical analysis was carried out using R (version 3.6.1) and RStudio (Version 1.2.5). Survival analysis and log-rank test were performed using the *survminer* and *survival* (Therneau, 2020) packages

## Acknowledgments

We would like to thank Julio Diaz Caballero and Jessica Hedge for their help in setting up the bioinformatic pipeline. This project was funded by Wellcome Trust Grant 106918/Z/15/Z held by RCM. We thank the Oxford Genomics Centre at the Wellcome Centre for Human Genetics (funded by Wellcome Trust grant reference 203141/Z/16/Z) for the generation and initial processing of the sequencing data. CS was supported by funding from the Biotechnology and Biological Sciences Research Council (BBSRC) [grant number BB/M011224/1]. JAE was supported by the European Research Council (ERC) through a Starting Grant (803375), the Atraccion de Talento Program of the Comunidad de Madrid (2016-T1/BIO-1105 and 2020-5A/BIO-19726) and the Ministerio de Ciencia e Innovacion (PID2020-117499RB-I00).

## Supplementary Figures

**Figure S1:**
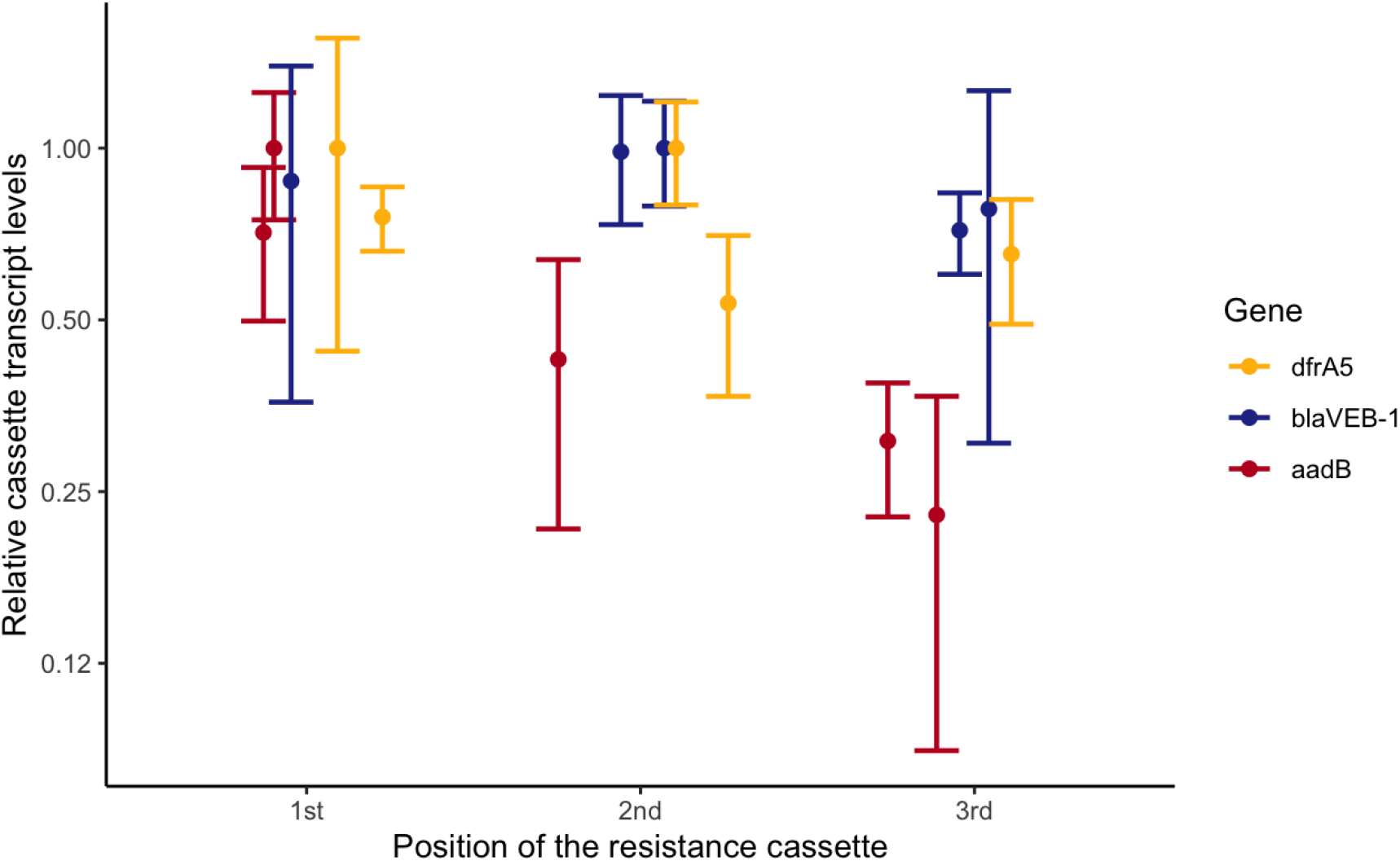
Transcript levels of each integron cassette depending on its position within the array relative to the cassette with the highest expression level. *aadB* expression data is reproduced from (Souque et al., 2021) and complemented with expression data for the two other cassettes. Cassette transcript levels are normalised based on two references genes and the plasmid copy number. Error bars represent the standard error of three independent biological replicates.

**Figure S2:**
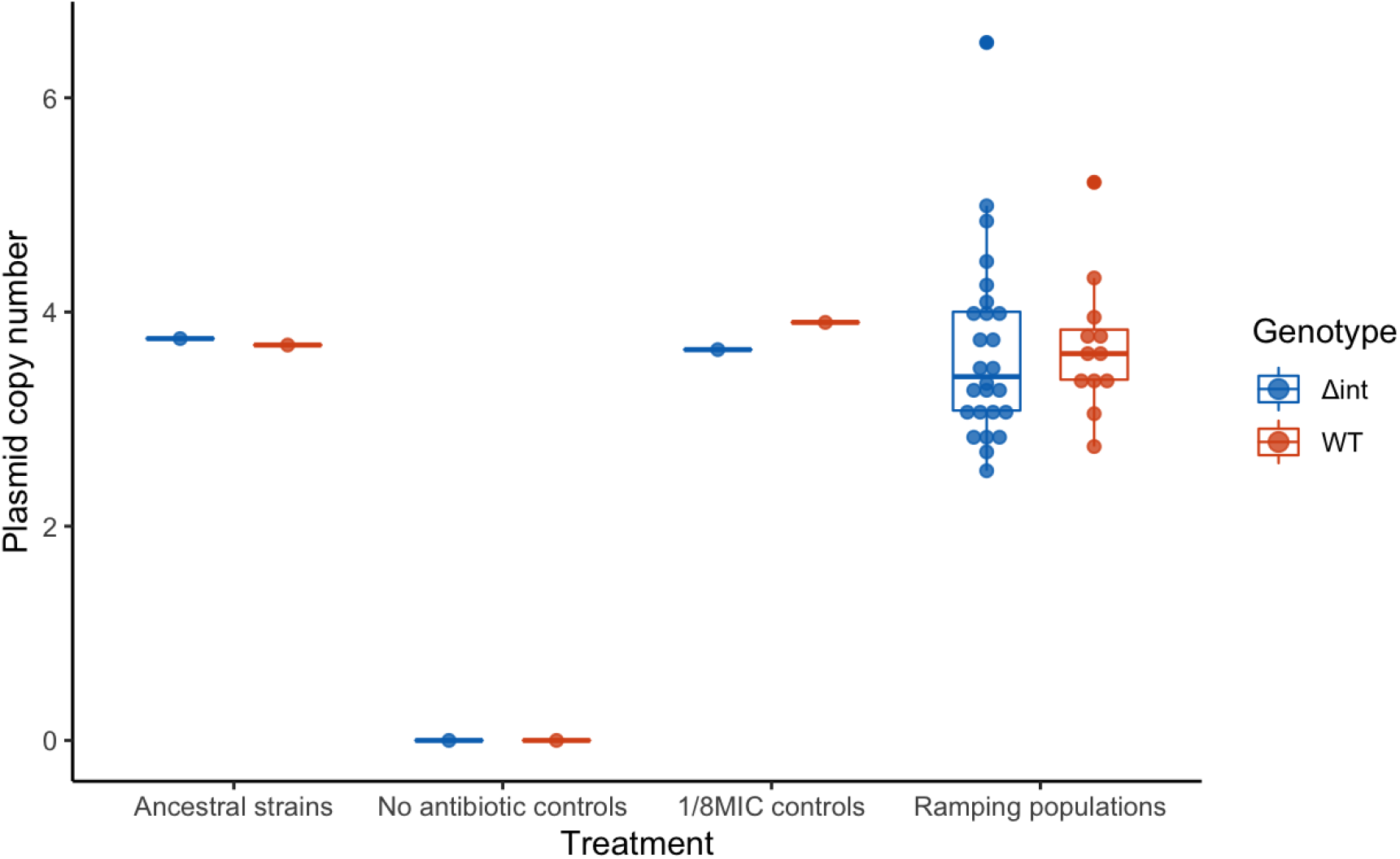
Plasmid copy number in evolved populations, estimated from sequencing data by dividing plasmid by chromosome average coverage. Control populations passaged either without antibiotic or at a constant concentration (1/8MIC) were mixed in an equimolar pool before sequencing.

**Figure S3:**
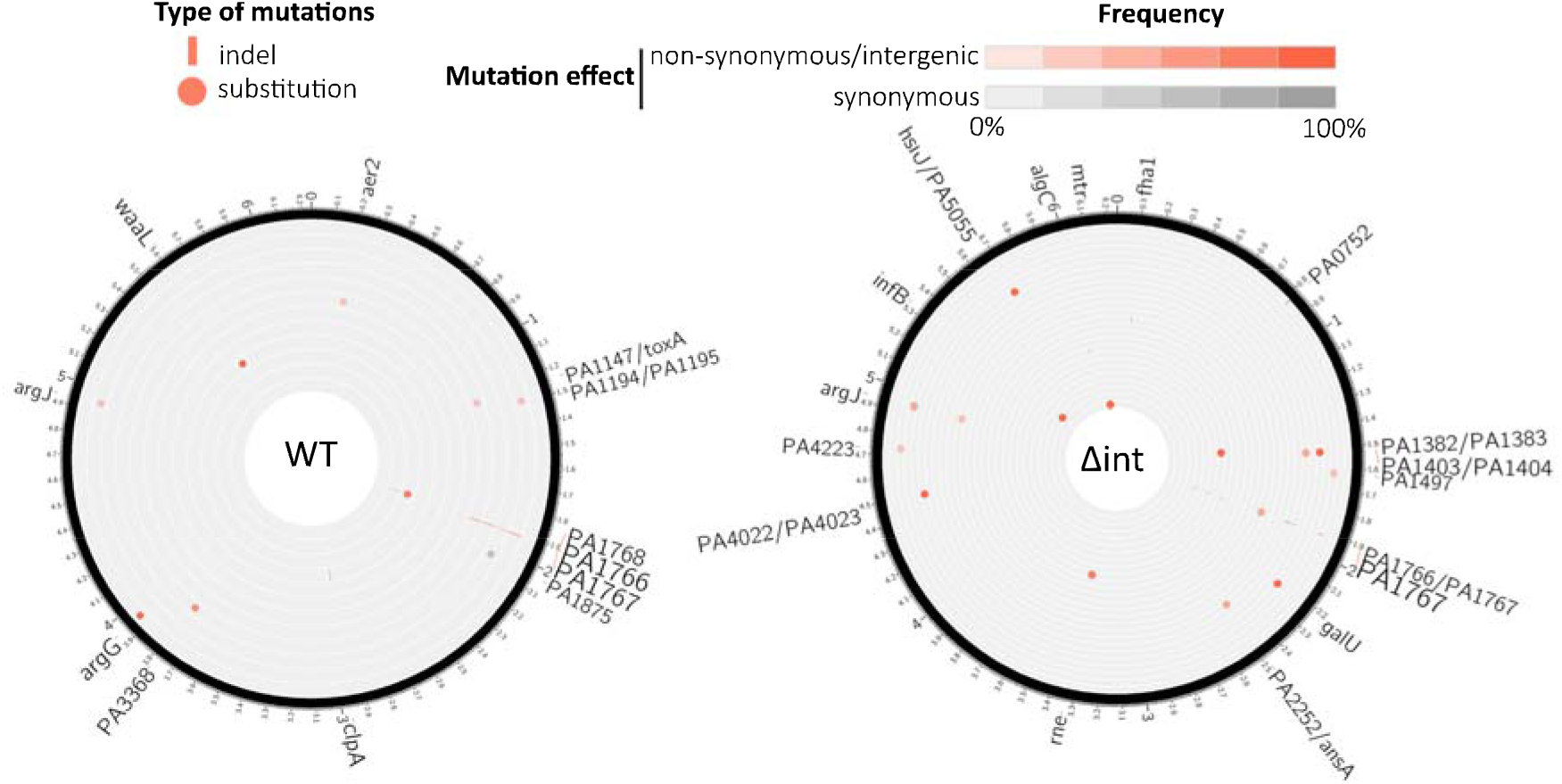
Chromosomal mutations in the surviving WT (left - 12 populations) and *Δint* (right - 26 populations) populations at x2 MIC mapped to the PA01 reference sequence. Each circle represents a separate population. The type (indel, substitution) of each mutation is represented by the shape of the marker (line, circle), while the marker colour represents the effect of the mutation (nonsynonymous/intergenic vs synonymous) and its colour intensity represents its frequency. The size of the gene labels on the outer ring represents the overall cumulative frequency of mutations present in this gene across all populations from this genotype.

## Supplementary Tables

- **S1: List of mutations and rearrangements identified by sequencing**
- **S2: List of strains & plasmids used in this study**
- **S3: List of primers**

## Author contributions

J.A.E. and R.C.M. conceived the project and J.A.E, C.S. and R.C.M conceived the experimental design. C.S. and J.A.E performed and analyzed the experiments. C.S. performed the bioinformatic analyses. C.S., J.A.E and R.C.M wrote the paper.

**The authors declare no competing financial interests**.

